# Fcγ-receptor-independent controlled activation of CD40 canonical signaling by novel therapeutic antibodies for cancer therapy

**DOI:** 10.1101/2023.01.16.521736

**Authors:** Carmen Reitinger, Karsten Beckmann, Anna Carle, Eva Blümle, Nicole Jurkschat, Claudia Paulmann, Sandra Prassl, Linda V. Kazandijan, Falk Nimmerjahn, Stephan Fischer

## Abstract

Activation of CD40-mediated signaling in antigen-presenting cells is a promising therapeutic strategy to promote immune responses against tumors. Agonistic anti-CD40 antibodies currently in development require Fcγ-receptor-mediated crosslinking of CD40 molecules for meaningful activation of CD40 signaling but have limitations due to dose-limiting toxicities. Here we describe the identification of CD40 antibodies which strongly stimulate antigen-presenting cells in an entirely Fc-independent manner. These novel Fc-silenced anti-CD40 antibodies induce upregulation of costimulatory receptors CD80 and CD86 and cytokine release by dendritic cells with an efficacy exceeding that of existing antibodies. Binding to the CD40L interaction region on CD40 appears to be a prerequisite to achieving such strong activities. Finally, the most active identified anti-CD40 antibody shows evidence of activity in terms of the expected markers of canonical CD40 signaling when injected in humanized mice. There are no signs of obvious toxicities whereas the clinical-stage anti-CD40 antibody CP-870,893 induced severe signs of toxicity in these animals despite a lower dose compared with the novel Fc-silenced canonical agonist. These studies thus demonstrate potent activation of antigen-presenting cells with anti-CD40 antibodies lacking Fcγ-receptor-binding activity and open the possibility of an efficacious and safe combination therapy for cancer patients.

**One Sentence Summary:** Treatment of antigen-presenting cells and humanized mice with novel Fc-silenced CD40 antibodies demonstrates an Fcγ-receptor-independent canonical agonistic mode of action for therapeutic use.

## Introduction

Over the past decades, a number of therapeutic strategies have been developed which are aimed at supporting cancer patients’ immune reaction to their tumor. Checkpoint inhibitors (CPIs) like PD1/PDL1-and CTLA-4-targeting antibodies have proven to be effective in several cancer types, yet a large fraction of patients whose tumor microenvironment is often classified as “cold” or non-inflamed do not respond to CPI therapy (*1*). In addition to testing combinations of different CPIs, recent approaches to tackling resistance to CPI therapy include CAR-T cell therapies or vaccinations with personalized neoantigens. However, such therapies are expensive; they require a large infrastructure and the therapeutic effects have been mixed.

The CD40-CD40L-signaling axis is a powerful, well-studied mechanism that is key for the generation of new, antigen-specific immune responses. Professional antigen-presenting cells expressing CD40 are stimulated by CD40L on T-helper cells in the course of antigen recognition. As a consequence, CD40 signaling licenses the APC for the stimulation of antigen-specific CD8 T-cells by the upregulation of costimulatory surface receptors and the release of T-cell-activating cytokines. The therapeutic triggering of canonical CD40 signaling is particularly attractive, since it provides an opportunity to elicit novel patient- and tumor-specific immune responses using a conventional biologic, e.g. an agonistic antibody. Experiments in mice have provided extensive support for this therapeutic strategy. In early studies FGK-45, an agonistic anti-murine-CD40 antibody, has been shown to stimulate cytotoxic T-cell responses and to provide protection against tumor re-challenge (*2*). Local intratumoral injection of FGK-45 further demonstrated systemic licensing of tumor-reactive cytotoxic T-cells resulting in the eradication of distant tumor nodules (*3*). This antigen-specific T-cell activation is the initially proposed primary signaling pathway controlled by CD40 activation (*4*).

The use of F(ab)2 antibody fragments and Fcγ-receptor-knock-out mice revealed that the activity of FGK-45 and other CD40 antibodies is largely mediated by Fcγ-receptor-directed crosslinking of CD40 molecules. In particular, FcγRIIb has been identified as a potent enhancer of anti-CD40 antibody-mediated activities (*5, 6*). For certain IgG2 Fc-variants an alternative CD40 crosslinking mechanism depending on the antibody conformation has been suggested (*7*), but overall, the process of Fcγ-mediated crosslinking has been established as the major mode of action for CD40 and other TNFR family-targeting agonistic antibodies (*8*). However, it was also recognized in many studies that treatment with CD40 antibodies is associated with severe side effects. For example, FGK-45, depending on the site and schedule of administration, elicited shock syndrome, liver damage, thromboembolic events and caused lethality (*3, 9, 10*). There is evidence indicating that the ability of CD40 antibodies to bind Fcγ-receptors provokes these adverse events. Mutations increasing CD40 antibody binding to Fcγ-receptors augmented thrombocytopenia, liver enzyme elevations, intravascular thrombosis, and liver damage (*11, 12*). Interestingly, the association of Fcγ-receptor binding and adverse events is not restricted to CD40 antibodies but has also been observed in antibodies targeting other TNFR family members or CD40L (*13–16*). These observations support the idea that alternative pathways are activated besides the initially defined canonical CD40 signaling leading to antigen-specific T-cell licensing.

Nevertheless, therapeutic CD40 antibodies entered clinical development more than ten years ago. The pharmacologic profile is reminiscent of what has been observed in animal models. The best-studied antibody CP-870,893 showed clinical activity but also severe side effects. Dose-limiting toxicities were thromboembolic events and liver enzyme elevations resulting in a maximum tolerated dose in patients of 0.2 mg/kg (*17–19*). Fc-modified second generation CD40 antibodies with increased binding specificity for FcγRIIb have been generated with the aim of achieving a better therapeutic window (*11, 20*). However, some of the Fc modifications aggravated the adverse events in preclinical in vivo models and intratumoral application has been proposed for clinical use (*11, 12*).

Based on this preclinical and clinical experience we reasoned that the Fcγ-receptor-dependent mode action of current CD40 antibodies is associated with a complex and problematic pharmacology which is due to non-canonical signaling events. The avoidance of Fcγ-receptor-binding capabilities appears reasonable but requires a different mode of action, whereby canonical CD40 signaling is efficiently triggered by crosslinking of CD40 via the paratope of the antibody but without the necessity for higher-level crosslinking via the Fc-part. The generation of such antibodies, however, is obviously a very rare event since such molecules have not been described so far. Therefore, we employed a highly efficient antibody discovery process to generate a maximum diversity of paratopes together with testing of large candidate numbers focused on the desired relevant function. In this study we describe the generation of novel CD40-targeting antibodies and their characterization in vitro and in vivo. We used immunization of wildtype rabbits combined with direct B-cell cloning, high-throughput functional screening and humanization. We received large numbers of diverse agonistic anti-CD40 antibody candidates. Careful functional selection and a precise focus on an Fcγ-receptor-independent mode of action resulted in humanized therapeutic antibodies with an extraordinary in vitro and in vivo activity profile.

## Results

### Identification of novel CD40 agonistic antibodies

The attractiveness of CD40 as a therapeutic target and the evident imbalance between safety and efficacy for current CD40-targeting therapeutics in development prompted us to identify novel antibodies which exhibit a significantly improved therapeutic window. Wildtype rabbits are a unique source of therapeutic antibodies because of their exceptional mechanisms for diversifying antibody gene sequences. Common VDJ recombination and somatic hypermutation mechanisms are enforced by V-gene conversion, which proceeds to diversify the antibody repertoire even during the course of an immunization, thus leading to diverse and high-affinity antibodies (*21, 22*).

We immunized rabbits with recombinant human CD40, isolated monoclonal B-cell clones by FACS and functionally tested > 16,000 antibodies secreted into the supernatants of cultured monoclonal B-cells (Figure 1A). The high-throughput HEK-Blue™ NFκB gene reporter screen identified more than 1,200 agonistic anti-CD40 antibody candidates. VL and VH genes of more than 300 antibody candidates were sequenced from B-cell lysates. We generated one humanized V-region sequence for each rabbit V-region sequence and cloned the humanized sequences into a human IgG1 backbone containing the L234A and L235A (LALA) mutation to prevent Fcγ-receptor binding (*23*). 198/303 candidates were found to be functional after humanization and recombinant production as hIgG1-LALA antibodies (Suppl. Fig. 1). Figure 1B shows the activity of 88 selected humanized anti-CD40 hIgG1-LALA antibodies in the HEK-Blue™ NFκB gene reporter assay. The data indicates that all 88 antibodies confer basic agonistic effects by binding to CD40.

**Figure 1:**
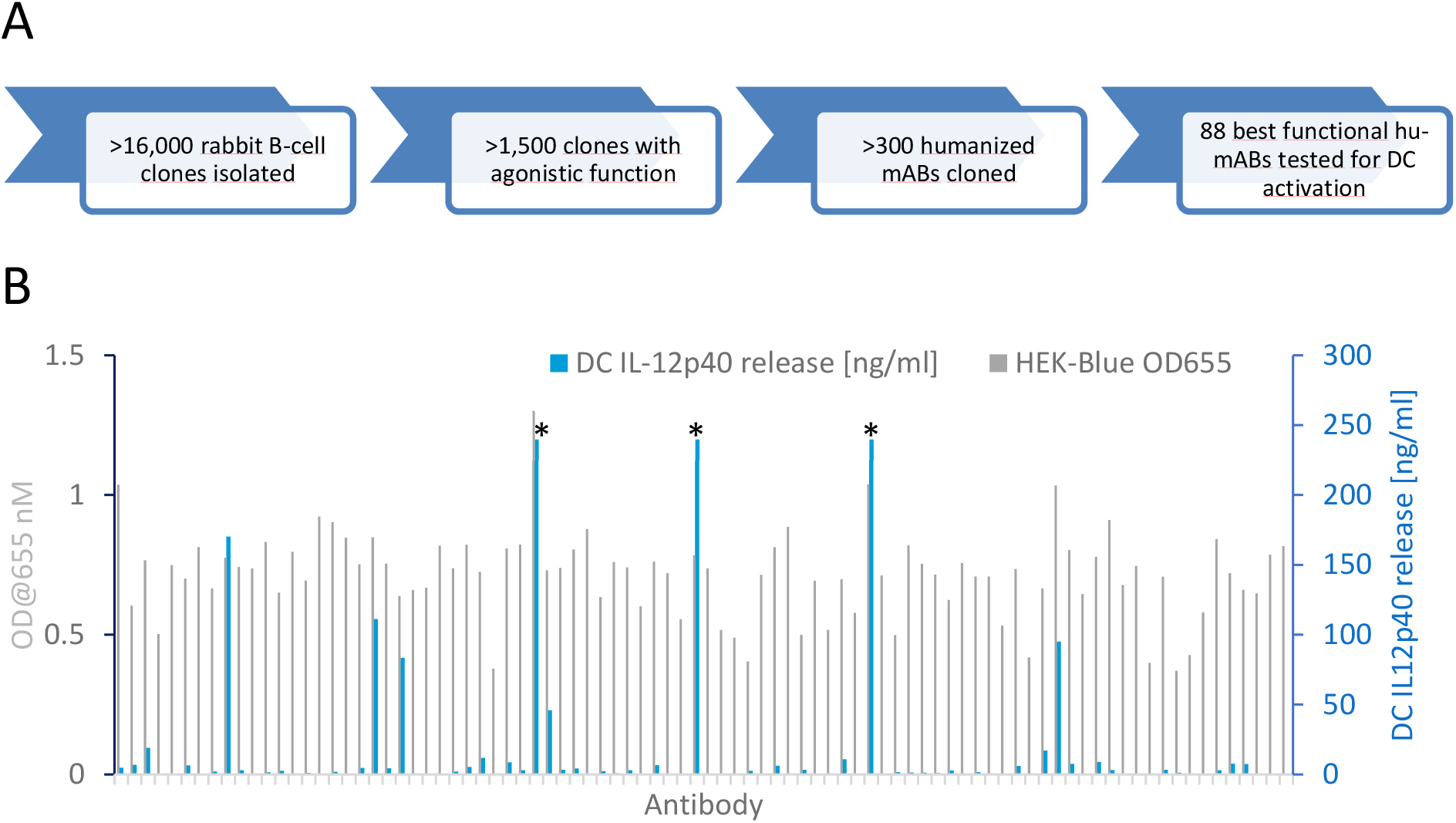
Identification of Fc-independent APC-activating anti-CD40 antibodies. (**A**) Flow diagram of the anti-CD40 antibody discovery campaign. Wildtype rabbits were immunized with recombinant CD40 protein and monoclonal B-cells were FACS-sorted and cultured. Antibodies secreted into the B-cell supernatant medium were screened for induction of NFκB signaling in a HEK-Blue-CD40L™ cell-based gene reporter assay. Antibody gene sequences were retrieved from B-cells, humanized and cloned as human IgG1-LALA antibodies in HEK-293 FreeStyle™ cells. (**B**) 88 humanized anti-CD40 hIgG1-LALA antibodies were tested in the HEK-Blue-CD40L™ cell-based gene reporter assay (grey bars; OD@655 = secreted embryonic alkaline phosphatase activity) and for the induction of IL-12p40 cytokine release by in vitro differentiated dendritic cells (blue bars; measured by ELISA). Asterisks indicate values exceeding the linear range of the ELISA.

### Fcγ-receptor-independent activation of innate immune responses

For further differentiation we established a primary cell-based, physiologically relevant test system. In vitro-differentiated DCs (moDCs) are known to express Fcγ-receptors (*24*) and stimulation of DC maturation is thus a suitable assay to test whether antibodies induce activation of CD40 signaling dependent on or independent of Fcγ-receptor-mediated crosslinking. Production and release of IL-12 by DCs is a hallmark of DC maturation which supports T-cell Th1 differentiation and antitumor effects and was used as a measure of efficacy (*25*). We analyzed the activity of the selected 88 antibodies and found to our surprise that a small fraction of the IgG1-LALA antibodies induced high levels of IL-12p40 release by moDCs, while the majority of antibodies show comparably low activity or no activities at all (Figure 1B).

We next tested whether the DC maturation assay was able to reflect Fcγ-receptor-dependent agonism, thus we compared the activities of the identified novel antibody LALA variants with Fc-variants of the clinical antibody CP-870,893. We generated the hIgG2 form, the clinical-stage antibody, and hIgG1, hIgG1-V11 and hIgG1-LALA variants. These molecules should provide different capabilities to interact with Fcγ-receptors (IgG1-V11 > IgG1 > IgG2 > IgG1-LALA). hIgG1-V11 contains mutations which increase the affinity for binding to FcγRIIb by 40-fold (*26*). In fact, CP-870,893-hIgG1-V11 and -hIgG1 show 12-fold or 5-fold increased activity while the hIgG1-LALA version shows about 3-fold reduced activity compared with the hIgG2 clinical antibody (Figure 2A). This experiment thus confirms that the impact of Fcγ-receptor binding on CP-870,893 activity can be determined in the assay. The four tested novel hIgG1-LALA antibodies 271, 273, 276 and 278 induced IL-12p40 release with activities 6- to 32-fold higher compared with CP-870,893-hIgG2 (Figure 2A). To rule out the possibility that the four antibodies harbor similar paratopes, we analyzed the antibody CDR-sequences to determine the degree of amino acid sequence divergence. As shown in Table S1, HC and LC CDRs differ by 20-58 %, indicating that these antibodies are not related to each other at the sequence level. In summary, these results provide convincing evidence that we were able to identify a diverse set of CD40-specific antibodies with the ability to induce a strong CD40-dependent activation of antigen-presenting cells independently of Fcγ-receptor-mediated crosslinking.

**Figure 2:**
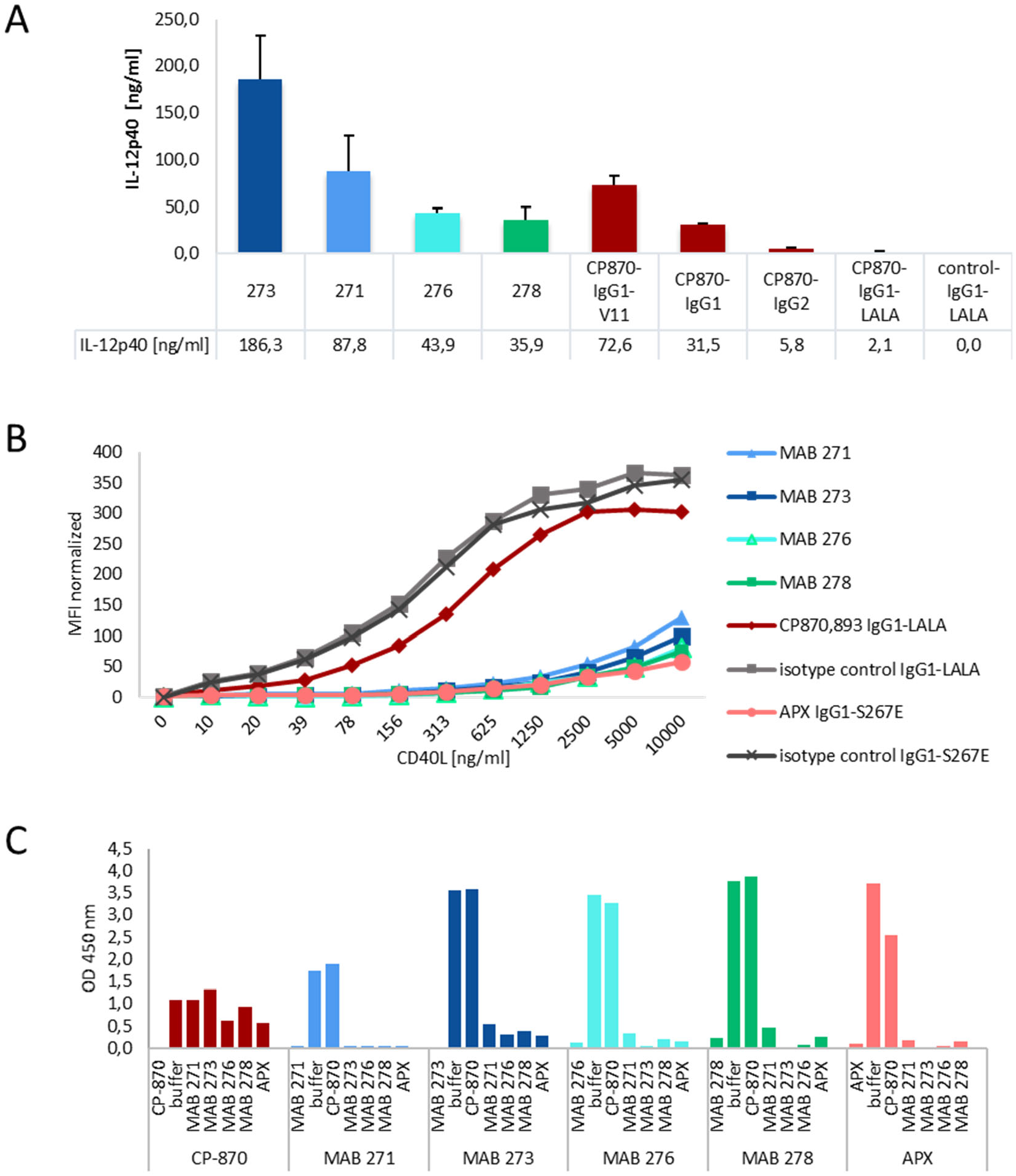
Strong activities on moDCs and binding to the CD40L binding region differentiate the novel Fcγ-receptor-independent anti-CD40 antibodies from CP-870,893. (**A**) The activity of the four most active CD40 hIgG1-LALA antibodies MAB 271, 273, 276, 278 (Figure 1B) were tested for the activation of in vitro differentiated DCs in direct comparison to different Fc-variants of CP-870,893. IL-12p40, released by DCs, was measured by ELISA. Error bars indicate StDev (mean) of triplicates. (**B**) CD40 antibodies were incubated with CD40-expressing HEK-Blue-CD40L™ cells and interference with CD40L binding to CD40 was tested by the addition of increasing concentrations of recombinant CD40L-mIgG2a-Fc-fusion protein. Binding of CD40L was detected by flowcytometry using a fluorescently labeled anti-mouse IgG antibody. (**C**) Competition of anti-CD40 antibodies for binding to human CD40 was tested in a sandwich ELISA. Each antibody was tested both as a coating and as a detection antibody in combination with any other anti-CD40 antibody.

### Agonistic antibody-mediated activation of CD40-dependent effector pathways

Next, we investigated whether these antibodies compete with the CD40 ligand binding to CD40. The aim of this experiment was to establish whether the four identified antibodies have a common mode of interaction with the CD40 receptor. As comparators we used CP-870,893, which is known not to bind the CD40L-binding region, and APX, a clinical-stage hIgG1 antibody with an S267E mutation in the Fc-part to increase FcγRIIb binding which does compete with CD40L (*20, 27, 28*). As shown in Figure 2B, all four identified MAB antibodies and APX blocked binding of increasing concentrations of mouse-Fc-tagged CD40L to CD40-expressing HEK-Blue™ cells. Isotype control and CP-870,893 antibodies did not interfere. In the same samples, we also tested binding of anti-CD40 antibodies to HEK-Blue™ cells in the presence of CD40L. Here, we detected the anti-CD40 antibodies by indirect fluorescence using a labeled anti-human-Fc antibody. As seen in suppl. Figure 2, CD40L did not significantly affect anti-CD40 antibody binding to these cells.

We then investigated the mutual CD40-binding competition of all antibodies using a sandwich ELISA setup where each antibody was tested either as a coating or as a detection antibody. In line with the results presented in Figure 2B, all CD40L-competing antibodies also competed with each other for binding to CD40, while none of these antibodies competed with CP-870,893 (Figure 2C). In summary, the data in Figure 2 show that the four novel CD40 antibodies analyzed confer Fcγ-receptor-independent stimulation of moDCs and that this activity appears to require binding to the CD40L interaction region on CD40.

Another hallmark of DC maturation is the upregulation of costimulatory receptors which support the MHC-restricted activation of CD4^+^ and CD8^+^ T-cells. Therefore, we investigated the dose-dependent upregulation of the major costimulatory receptors CD86 and CD80 and the MHC molecule HLA-DR by flow cytometry of moDCs treated with the two most active novel anti-CD40 antibodies, MAB 271 and MAB 273. As comparator antibodies we included CP-870,893-IgG2 and APX-hIgG1-S267E to cover both CD40L-competing and non-competing Fcγ-receptor-dependent anti-CD40 antibodies. The results in Figure 3A show that all antibodies induce coreceptor expression in a dose-dependent manner. MAB 273 and 271 treatment induced the highest expression levels of the coreceptors tested while the maximum activities of CP-870,893 and APX were about two- to four-fold lower (Figure 3A and B).

**Figure 3:**
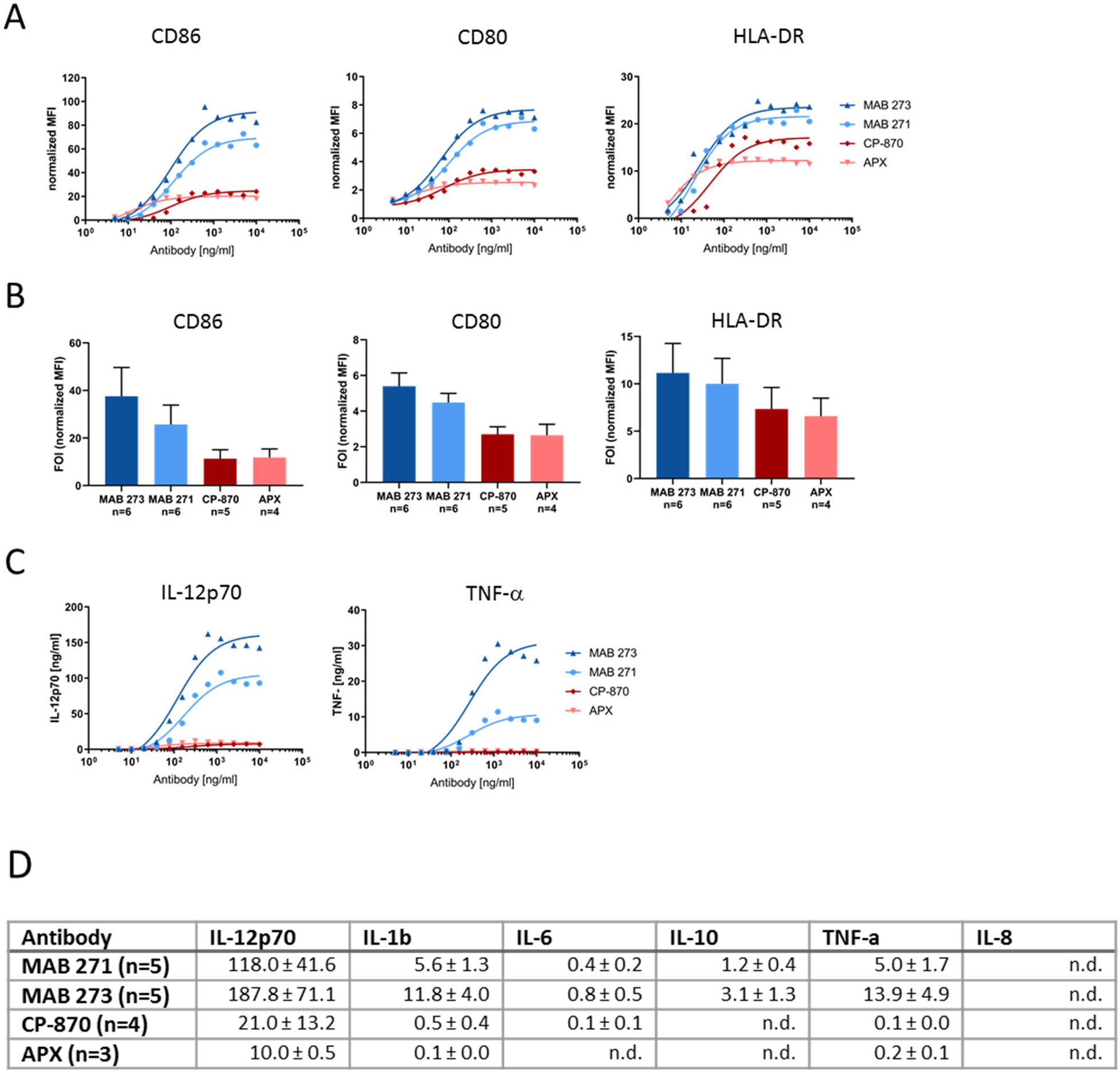
Induction of costimulatory receptors and cytokine release by Fc-silenced anti-CD40 antibodies in dendritic cells. Human PBMC-derived moDCs were treated for 2 days with anti-CD40 hIgG1-LALA antibodies MAB 271 and MAB 273, CP-870,893 hIgG2, APX hIgG1-S276E or isotype control antibodies. Expression of CD80, CD86 and HLA-DR was measured by flow cytometry (**A, B**) and cytokine concentrations in culture supernatants were determined by a cytometric bead array (**C, D**). MFIs were normalized to isotype control antibody treatments. In A and C a dose response analysis with DCs of an exemplary donor is shown, (**B**) and (**D**) summarize the mean of indicated donors treated with saturating concentrations of 1.25 or 2 μg/ml. Errors (**D**) and error bars (**B**) show SEM. n.d. = not detectable.

Next, we characterized the inflammatory cytokine secretion profile of anti-CD40-treated moDCs using a proinflammatory cytokine bead array. Mature IL-12p70, which is driving Th1 differentiation, is the cytokine most significantly induced by MAB 273 and 271, while the Th2 cytokine IL-10 is secreted at low levels (Figure 3C and D). Compared with MAB 273, CP-870,893 and APX induced about a 9- and an 18-fold lower IL12p70 cytokine release, respectively. Furthermore, TNF-alpha and IL-1β, both known to stimulate or to be produced by DCs undergoing maturation (*29*), are induced at least 20-fold higher with MAB 273 compared with CP-870,893- and APX-treated moDCs (Figure 3C,D).

Besides professional antigen-presenting DCs, CD40 is also expressed on B-cells. Like DCs, B-cells respond to CD40 activation with upregulation of CD80 and CD86 to increase antigen-presenting capabilities (*30, 31*). Therefore, we next determined the activity of the anti-CD40 antibodies to induce upregulation of these costimulatory receptors on B-cells isolated from human PBMCs from different donors. Compared with DC activation (Figure 3), we observed an overall less pronounced upregulation of CD80 and CD86 induced by the antibodies (Figure 4). As observed in moDCs, all antibodies induced CD80 and CD86 expression in a dose-dependent manner (Figure 4A). However, the upregulation of both receptors varied significantly from donor to donor. Overall, the mean activities of MAB 273, CP-870,893 and APX in four different donors are similar while coreceptor induction induced by MAB 271 appears to be relatively lower (Figure 4B). Taken together, the in vitro data demonstrate that canonical CD40 signaling, leading to cytokine release and costimulatory receptor upregulation on primary DCs and B-cells, is strongly induced by the novel Fc-silenced anti-CD40 antibodies.

**Figure 4:**
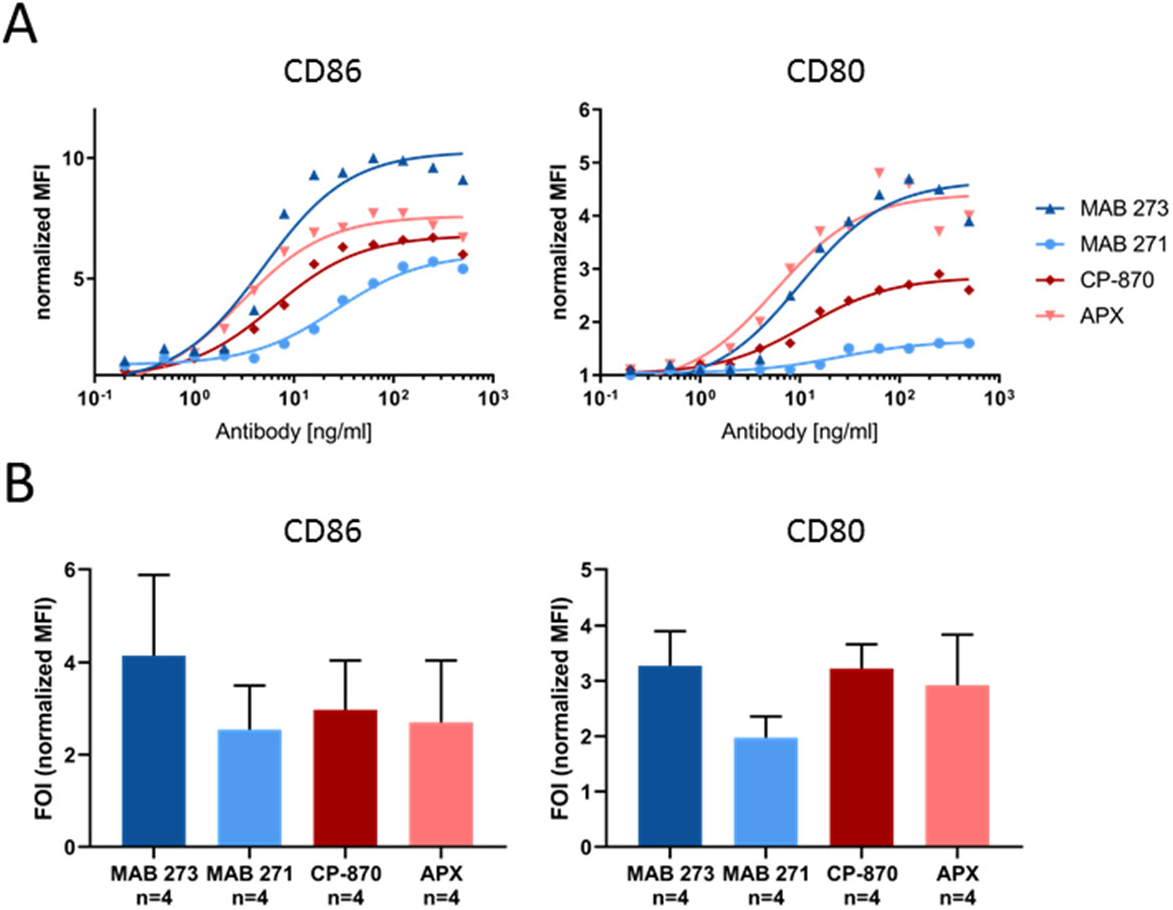
Fc-silenced anti-CD40 antibodies induce costimulatory receptors on human B-cells. B-cells were isolated from human PBMCs and incubated with anti-CD40 hIgG1-LALA antibodies MAB 271 and MAB 273, CP-870,893-hIgG2, APX hIgG1-S276E or isotype control antibodies. Expression of CD80, CD86 was measured by flow cytometry. MFIs were normalized to isotype control antibody treatments. In (**A**), a dose response analysis with B-cells of an exemplary donor is shown, (**B**) summarizes the mean of four donors treated with saturating concentrations of 312 μg/ml (1 donor) or 500 μg/ml (3 donors). Error bars indicate SEM.

### Assessment of pharmacodynamic markers and potential side effects of the CD40 agonistic antibody MAB 273 in humanized mice

The preclinical and clinical experience with CD40-targeting therapeutic antibodies has shown proof of concept but has also demonstrated substantial limitation of current therapeutic molecules due to toxicities. Thromboembolic events and liver transaminase elevations have resulted in a maximum tolerated dose of only 0.2 mg/kg for CP-870,893 (*17–19*). Therefore, an important question is whether the Fcγ-receptor-independent mode of action translates into a relevant biological activity in vivo and whether treatment is associated with potential side effects. The most efficacious novel agonistic CD40-specific antibody clone MAB 273 was tested in vivo. As a model we made use of human-stem-cell-transplanted mice with an established human immune system and which lack murine Fcγ-receptor expression by genetic deletion (*32*). These mice allow the study of anti-human-CD40 antibody function and potential side effects, considering contributions of human Fcγ-receptor but avoiding interference of murine Fcγ-receptors expressed on mouse-innate immune effector cells. For comparison the clinical hIgG2 antibody CP-870,893 was used. A single injection of 3 mg/kg of this antibody resulted in toxicity evidenced by a loss of body weight and a drop in body temperature on day 3 as shown in figure 5. Across independent experiments, 5 out of all 13 mice which had been treated with 3 mg/kg CP-870,893 had to be sacrificed in response to treatment (Figure 5 and Table S1). In contrast, treatment with 10 mg/kg MAB 273 was tolerated well with no evident drop in body weight or temperature over the course of 10 days (Figure 5). Histological analyses of kidney and liver revealed no obvious signs of tissue damage or inflammation for either the MAB 273 or CP-870,893 antibodies. No abnormalities were detected in HE staining of kidneys and livers in any animal. However, in three CP-870,893-treated mice immunofluorescent staining analysis showed murine platelet, and, in one mouse also murine macrophage infiltration in the liver (suppl. Figure 3). There was a detectable increase in T-cell numbers in the livers of all MAB 273-treated mice but not in isotype control hIgG1-LALA- or CP-870,893-treated mice (suppl. Figure 3). Liver transaminase levels in serum have not been elevated in any treatment group (suppl. Figure 4). In summary, MAB 273 treatment is tolerated by the humanized mice at the tested dose of 10 mg/kg, whereas treatment with 3 mg/kg CP-870,893 induced severe side effects thus precluding treatment at higher doses.

**Figure 5:**
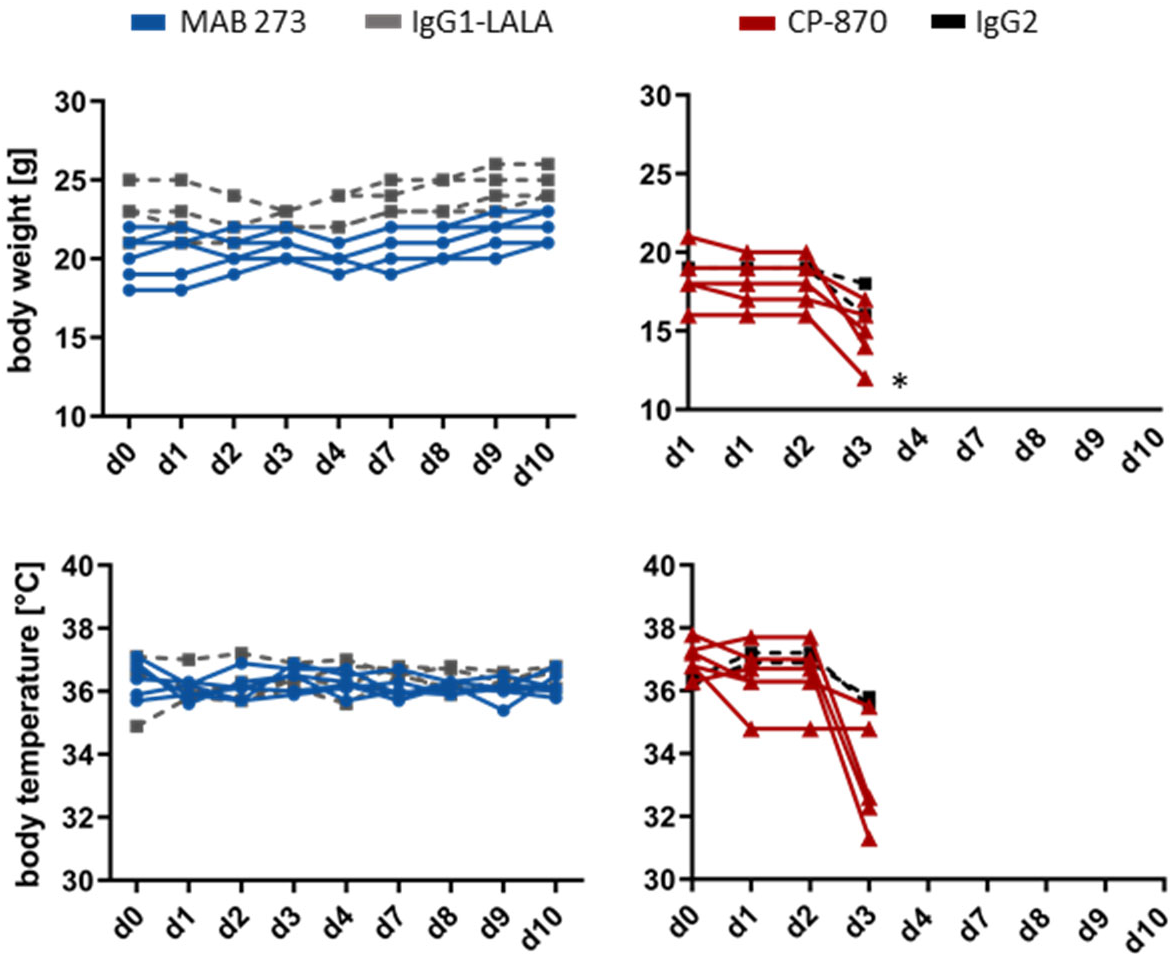
Changes in body weight and body temperature of human-stem-cell-transplanted mice after anti-CD40 antibody injection. Antibodies were injected on day 0 and measurement was done on different days after treatment. Injection of 6 mice each with MAB 273 (10mg/kg) and CP-870,893 (3mg/kg) was performed in two experimental runs separate from one another. Duration of the experimental run was 10 days for MAB 273 and 3 days for CP-870,893. *Data points of two mice are superposed.

We also tested whether there is evidence for on-target efficacy in the antibody-treated mice. Therefore, in a cohort of mice treated with MAB 273 (10 mg/kg), CP-870,893 (3 mg/kg) or hIgG1-LALA (10 mg/kg), we analyzed the development of blood cell numbers and concentrations of cytokines in the plasma over the course of 10 days after a single antibody injection (Figure 6). One out of four mice treated with CP-870,893 in this cohort had to be sacrificed on day 4. Confirming observations in other preclinical and clinical studies, we found an immediate disappearance of circulating B-cell in the blood in CP-870,893- and MAB 273-treated mice (Figure 6A). The numbers of NK- and T-cells appear to increase significantly until day 10 in MAB 273-treated mice but remain stable in CP-870,893-treated mice (Figure 6A). Importantly, proinflammatory cytokines like IL12p70, IL-6, TNF-a and IFN-γ, promoting or being produced as a result of Th1 differentiation and proliferation of T- and NK-cells, are also increased in the serum of MAB 273-but not CP-870,893-treated mice (Figure 6B). These in vivo results are in line with the relative activities of MAB 273 and CP-870,893 measured in vitro in moDCs and provide evidence that cells of the adaptive immune system are being activated in response to a single dose of MAB 273. All in all, this study supports a new paradigm of therapeutic targeting of the CD40 receptor. In addition, the study provides evidence that treatment with an anti-CD40 antibody that is independent of Fcγ-receptor crosslinking does not induce evident toxicities in the applied model.

**Figure 6:**
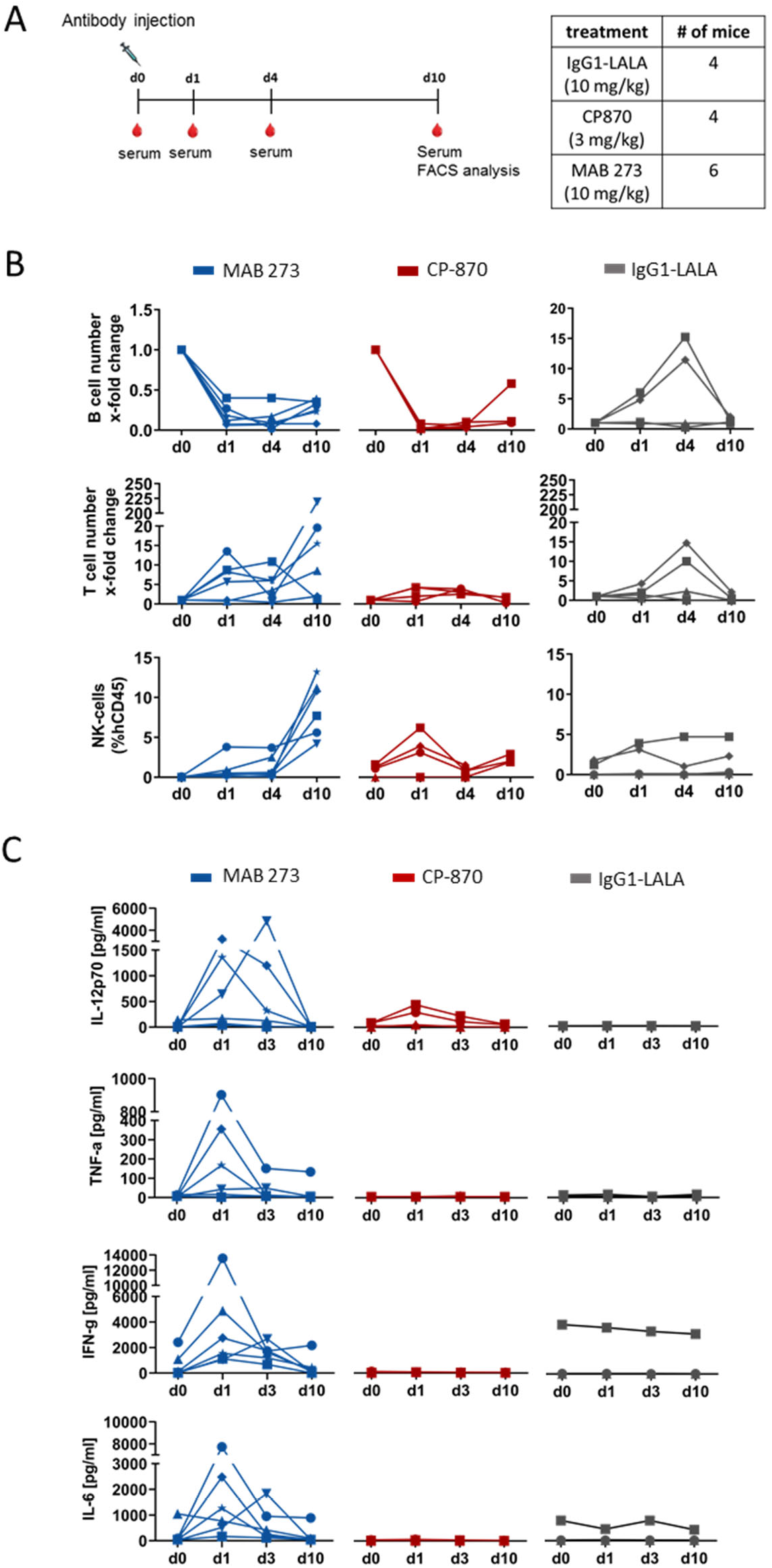
Effect of anti-CD40 antibody injection on human immune cell numbers and cytokine production in a human-stem-cell-transplanted mouse model. (**A**) Experimental setup. The table on the right shows the number of mice and analyzed antibodies included in the presented experiment. (**B**) Analysis of peripheral blood leucocytes was performed by flow cytometry and alteration of the human B-cell and T-cell number is represented as x-fold change on the indicated time points relative to day 0 for each mouse treated with the respective antibodies. Human NK cell numbers (CD56+ CD16+ subpopulation) are shown in per cent of total human CD45+ cells in the peripheral blood of mice treated with the respective antibodies at the indicated time points. Symbols indicate individual mice. (**C**) Cytokine concentrations in murine serum were determined using a LEGENDplex™ MultiAnalyte Flow Assay. The graphs depict the concentration of human IL-12p70, TNF-α, IL-6 and IFN-γ on the indicated time points after injection of MAB 273, CP-870,893 and IgG1-LALA control antibodies. Symbols indicate individual mice.

## Discussion

The activation of tumor-specific immune responses in cancer patients through the use of agonistic and checkpoint control antibodies has become one of the most successful advances in cancer therapy over the past decade. Anti-PD1/PDL1- and -CTLA-4-blocking antibodies have proven to be efficacious in many cancers by “releasing the brakes” of ongoing but suppressed immune responses. However, since many patients do not respond to such therapies, there are numerous alternative and additional compounds in development. There is the expectation that, in combination with CPI therapy, a dramatic improvement of response rates should be possible (*33*). In fact, one of the major reasons for patients not responding to CPI therapy is the lack of pre-existing tumor-specific T-cells which CPI therapy could support (*34*). Therefore, combination therapies are warranted which promote the de-novo response to tumor antigens. Patient-specific vaccination strategies or adjuvant chemo- or radiotherapy are possible interventions to make tumor-derived neoantigens accessible for the immune system. It appears, however, that the efficient presentation of these neoantigens to tumor-specific CD4^+^ and CD8^+^ T-cells and hence their activation and differentiation into T-helper and cytotoxic T-cells is a bottle neck for the initiation of an efficacious antitumor response (*35*).

The triggering of CD40 receptor signaling on professional antigen-presenting cells has been shown to be a powerful way of supporting neoantigen presentation to T-cells in a great number of preclinical studies and clinical proof of concept has been demonstrated (*36*). Mechanistically, all CD40-targeting therapeutic biologics currently in development are based on higher-level crosslinking mechanisms. In the case of agonistic antibodies, Fcγ-receptor binding by accessory cells promotes CD40 crosslinking on APCs (*8*). Regarding the use of soluble CD40L as therapeutic agonists it should be noted that simultaneous interaction of CD40L with CD40 and multiple integrins has been shown which may also promote CD40 clustering (*8, 37*). It is worth noting that one of these integrins is the platelet-specific αIIbβ3 and its interaction with CD40L promotes thrombus stability (*38*). Blocking IgG1 antibodies targeting CD40L for the therapy of autoimmune diseases has also been shown to cause cardiovascular thrombotic events in clinical studies (*39*). Importantly, the avoidance of Fcγ-receptor interactions by using either an anti-CD40L Fab’ or an a-glycosylated IgG resulted in the disappearance of such side effects in a preclinical study (*13*). These examples indicate that CD40L pathway-targeting therapeutic molecules allowing higher-level crosslinking can cause significant side effects. For agonistic, Fcγ-receptor-binding CD40 antibodies which allow concomitant CD40L binding, the possibilities of molecular interactions and participating cell types are becoming even more complex. In fact, synergistic effects of a CD40L-non-competing anti-CD40 antibody with CD40L function have been emphasized as a way to enhance therapeutic efficacy (*40*). We see two problematic consequences associated with this concept. First, the clinical pharmacology is probably too difficult to control. Independent variables such as patient-specific expression and distribution of Fcγ-receptors, the induction of CD40L and the release of soluble CD40L can play a relevant role. Second, the capability to form multimeric complexes and the induction of costimulatory signals through crosslinking of cells and platelets harbors the risk of an uncontrolled pharmacology and of causing toxicities. Supporting this notion, Fcγ-receptor interactions have been shown to contribute to toxicities induced by antibodies targeting CD40 and other TNFR family members (*11,12, 15, 16*). This indicates that the ability to crosslink via Fcγ-receptors is a general liability of TNFR-targeting antibodies. In fact, vascular thromboembolic events and liver toxicities have been observed in patients treated with the hIgG2 anti-CD40 antibody CP-870,893 and resulted in an MTD of only 0.2 mg/kg (*17*).

Recently, it has been suggested that “second generation” CD40 agonists, furnished with optimized Fcγ-receptor-binding capabilities, will provide better clinical activity profiles (*36*). For example, the use of alternative isotypes and the introduction of specific Fc-mutations with increased Fc-receptor affinities and selectivity, in particular to the FcγRIIb receptor with CP-870,893, have been shown to boost the antibodies’ activities in vivo. However, at the same time toxicities increased too, and these could only be reduced by using an intratumoral route of antibody injection (*7, 11, 12*). Our results are in perfect agreement with these reports. The activation of DCs by CP-870,893 shows a strong dependency on the Fc-variants’ capability to interact with Fcγ-receptors (Figure 2A). Richman et al have shown that the activity of CP-870,893 is predominantly Fc-independent and they conclude that the epitope recognized on the CD40 molecule dictates the activity of this antibody (*6*). They compared the IgG2 form of CP-870,893 with a F(ab)2’ form. In agreement with those results, we observe basal activity of the IgG1-LALA version of CP-870,893 in activating DCs. However, it is important to note that the activity of this antibody is strikingly increased when the antibody is furnished with Fc-parts capable of efficient Fcγ-receptor-mediated crosslinking (Figure 2A).

Here, we show that we have isolated and characterized CD40 antibodies, which efficiently activate CD40-mediated signaling in a way that is solely dependent on the paratope-epitope interaction. In fact, the activities are higher compared with Fcγ-receptor optimized versions of CP-870,893 (Figure 3A). It is important to note here that the identification of such antibodies appears to be a rare event. Out of more than 1,500 CD40-activating candidate antibodies, only a small number displayed strong Fcγ-receptor-independent activation of DCs. This demonstrates the highly diverse antibody repertoire that rabbits are able to generate and justifies the requirement for testing large numbers of antibodies to identify such unique functionalities.

Regarding functional epitopes on CD40, a recent study comparing a number of anti-CD40 antibodies binding to different sites on CD40 concluded that antibodies binding to extracellular domains comprising the CD40L-binding region are less potent agonists and that the N-terminal domain is the region of choice to elicit potent agonism (*27*). In contrast, we find that binding to the CD40L-interacting region on CD40 appears to be a typical feature in connection with the strong activities of the novel antibodies identified in this study. The top four antibodies all interfere with CD40L binding to CD40 and they bind to the receptor in a mutually exclusive fashion (Figure 2B,C). In fact, we found even more antibodies binding to that region and these comprise functional as well as completely non-functional antibodies as determined by moDC IL-12 release. Strikingly, the analysis of heavy- and light-chain CDR3 sequences of the top four antibodies shows that the clones are not just unrelated, but very diverse and thus most probably bind in very different ways. Therefore, while binding to the CD40L region appears to be a prerequisite for the Fcγ-receptor-independent activity, the paratope of the antibody and not the epitope is the critical determinant for triggering canonical CD40 signaling in this study. For TNF family receptors it is assumed that the geometry of receptor clusters at the cell surface plays an important role in the efficiency of signal transmission (*41*). For CD40 the presence of the homodimeric form of the receptor on the cell surface has been linked to receptor function (*42, 43*). Thus, certain paratope-epitope interactions may induce antibody-CD40-dimer complex geometries at the cell surface which are particularly potent in transmitting signals across the cell membrane.

The identified MAB 273 antibody displays highly superior efficacy in vitro when compared with both the Fc-optimized CP-870,893-hIgG1-V11 and the CD40L-competing Fc-optimized hIgG1 APX antibodies (Figure 2A and 3). We are not aware of any CD40-targeting biologic, neither a recombinant CD40L nor an antibody, which produced effects in APCs that differentiate from these two comparator antibodies in a similar fashion. The differentiation regarding the induction of costimulatory receptors of MAB 273 on a panel on human B-cells was less prominent. One explanation for this may be the differential expression of Fcγ-receptors on different cell types. Whereas DCs express FcγRIIa, FcγRIIb and FcγRI, B-cells exclusively express the FcγRIIb and thus may optimally promote Fc-dependent agonistic anti-CD40 antibodies in the homogenous B-cell assay. It is worthy of note that the contribution of B-cells to therapeutic efficacy has been questioned recently, since in a syngeneic pancreatic cancer model B-cells were found not to be required for the observed effects using anti-mouse-CD40 antibody FGK-45 in vivo (*44*).

We aimed in this study to test our hypothesis that potent activation of CD40 signaling without Fcγ-receptor-mediated crosslinking is possible and may provoke fewer side effects in vivo. In order to compare our most effective antibody MAB 273 with CP-870,893, the best-studied clinically anti-CD40 antibody, it was necessary to choose a humanized mouse model in which antibody binding to human Fcγ-receptors is provided without the disturbance of mouse Fcγ-receptor interactions. The murine Fcγ-receptor-deficient human-stem-cell-transplanted mice used in this study constitute a unique model, shown to be well-suited to the study of Fc contributions of human IgG therapeutic antibodies (*32, 45*). Strikingly, a single injection of 3 mg/kg CP-870,893 resulted in obvious toxicity and 5 animals in a total of 13 treated animals had to be sacrificed. This is in line with the fact that in patients CP-870,893 is tolerated only at low doses with DLTs becoming apparent at 0.3 mg/kg. In particular, D-dimer elevations and thromboembolisms have been observed, indicating effects on platelet activation and coagulation. In our study it remains unclear what causes the fatal outcomes in CP-870,893-treated mice. Platelet infiltrates in liver tissue have been observed in some animals but in all other analyzed mice liver and kidney histology looked normal. Coagulative necrosis in the liver, as it has been shown to occur in mice treated with anti-murine-CD40 antibody FGK-45, could not be observed (*9, 46*). However, it cannot be ruled out that microvascular occlusions in other organs occurred in those animals. It is evident that the toxicity of the treatment is not related to cytokine release. In fact, CP-870,893 does not induce significant cytokine levels in the blood plasma of treated mice at the tested dose. In contrast, MAB 273 treatment leads to significantly increased blood plasma levels of proinflammatory cytokines such as IL-12, TNF-alpha, IL-6 and IFN-γ. This reflects perfectly the observations made in vitro, where IL-12 and TNF-alpha release by moDCs treated with MAB 273 was reproducibly demonstrated (Figure 3). These cytokines appear also to be responsible for the relative increases in T-cell and NK-cell counts which were observed subsequent to cytokine peak levels in MAB 273-but not CP-870,893-treated animals (Figure 3).

Another interesting observation is that CP-870,893 treatment strongly reduced B-cell numbers as it has been described in clinical studies. The model thus confirms the clinical effects of CP-870,893 concerning the occurrence of toxicities and B-cell reductions. The disappearance of B-cells has been interpreted as a pharmacodynamic marker in clinical studies, but it is not clear whether this is due to redistribution or cell depletion. In our in vivo model the effect on B-cells does not correlate with blood cytokine levels. In both MAB 273- and CP-870,893-treated animals, peripheral B-cell counts are significantly reduced but cytokines are induced only by MAB 273. This supports the idea that CP-870,893 binding to CD40 has qualitatively different functional consequences when compared with MAB 273 binding to CD40 in vivo. Published data show that Fcγ-receptor interactions of CD40-targeting antibodies strongly contribute to toxicities in preclinical studies (*11*). There, the authors concluded that efficacy and potential toxicity cannot be separated when an agonistic antibody is used. Currently, second-generation anti-CD40 antibodies are underway with engineered Fc-parts for optimal Fcγ-receptor involvement (*20*). Furthermore, preclinical studies have suggested the intratumoral injection of agonistic anti-CD40 antibodies to manage the intrinsic compound toxicities. This concept is currently being tested in clinical studies (*11, 47, 48*). However, our study and the functional profile of MAB 273 enable a new view of how the CD40 receptor system could be employed therapeutically. We show here that a strong CD40 signal induction on APCs is possible with anti-CD40 antibodies lacking Fcγ-receptor interaction capabilities. In addition, we provide preliminary evidence in a humanized mouse model, that therapy employing such antibodies may have a significantly improved therapeutic window. Importantly, toxicology studies for such antibodies in non-human primates are expected to provide predictive data since the toxicology profile will not be impacted by the differential Fcγ-receptor-binding capacities between humans and non-human primates (*49*). Providing an effective drug without adding toxicities is highly relevant for later combination studies. In summary, the anti-CD40 hIgG1-LALA antibodies identified constitute a highly efficacious, entirely novel agonistic CD40-targeting concept that is solely dependent on the antibody’s paratope interaction with CD40 and may significantly enhance cancer therapy as a single agent or in combination with established treatments.

## Material and Methods

### Antibodies and reagents

CD86-VioBright-515, CD80-PE and HLA-DR-VioBlue antibodies and isotype controls were from Miltenyi Biotec. Recombinant CD40L protein, containing a mouse-IgG2a Fc-tag was from AB Biosciences (#P7005F). HIS-tagged CD40 recombinant protein was from Acro Biosystems. Human-IgG2, IgG1, IgG1-V11 and IgG1-LALA antibody variants of CP870,893 contain the variable region sequences of the clone 21.4.1 described in patent US 7,338,660. The IgG1-V11 mutation is described in (*26*). APX is a hIgG1-S267E anti-CD40 antibody containing the variable region sequences of the antibody APX005 described in patent US 2014/0120103. MAB 273, 271, 276 and 278 antibodies were identified and cloned from human-CD40-immunized New Zealand white rabbit B-cells. Humanization was done at Fusion Antibodies plc. Variable regions were fused to hIgG1 Fc sequences containing L234A and L235A mutations (*23*). Antibodies were produced at MAB Discovery GmbH using the FreeStyle™ 293 expression system from Thermo Fisher. Transfected HEK-293 FreeStyle™ cells were incubated for 10-11 days at 37°C. Harvesting of cell supernatants was done by a 2-step centrifugation procedure, followed by a filtration step. Antibody purification from cell supernatants was done in two steps using the ÄKTA Avant purification system (GE Healthcare). The antibodies were purified by affinity chromatography using a Protein A resin (MabSelect SuRe, GE Healthcare), followed by a preparative size exclusion chromatography (SEC). All preparations were endotoxin-low (< 0,05 EU/mg) as determined by the EndoZyme^®^ assay (Hyglos GmbH, Germany).

### HEK-Blue-CD40L^TM^ cell-based gene reporter assay

The agonistic activity of anti-CD40 monoclonal antibodies was tested by stimulating HEK-Blue-CD40L™ (Invivogen) cells which harbor an NF-κB-inducible Secreted Embryonic Alkaline Phosphatase (SEAP) gene construct. 25,000 HEK-Blue-CD40L™ cells/well in 20μl DMEM containing 10% FBS were seeded in a cell-culture 384-well plate and cultured overnight. Humanized anti-CD40 hIgG1-LALA antibodies were added in a volume of 5μl medium to final concentrations ranging from 10 to 0.08 μg/ml. After 6 hours of incubation at 37°C and 5% CO_2_, 5μl of the medium supernatant of each well was transferred to a white clear-bottom 384-well plate containing 20μl of 2x QUANTI-Blue™ reagent (InvivoGen). After incubation at 37°C and 5% CO_2_ for one hour, optical density at a wavelength of 655 nM was measured reflecting NF-κB-dependent activation of phosphatase secretion.

### Induction of costimulatory receptors and cytokine release in dendritic cells

Buffy coats from different donors were provided by the Bavarian Red Cross. Monocytes were isolated from buffy coat-derived PBMCs by CD14 MACS (Miltenyi) according to the supplier’s protocol and differentiated to immature DCs (iDCs) for 5 days in the presence of 50 ng/ml hGM-CSF and 10 ng/ml hIL-4. Anti-CD40 or isotype control antibodies were added to iDCs in a 96-well plate (10^6^ cells/ml) at concentrations ranging from 10,000 to 5 ng/ml. For analysis of costimulatory receptors, DCs were harvested 48h after the addition of antibodies and stained using fluorophore-labeled antibodies against HLA-DR, CD86 and CD80. Analysis was performed by flow cytometry on a BD FACSVerse device. Cytokine concentrations in the supernatants of DC cultures were measured using a BD human inflammatory cytometric bead array kit (BD #551811) according to the manufacturer’s instructions. IL-12p40 cytokine release was quantified using the Duo Set ELISA kit from R&D Systems according to the manufacturer’s instructions. Absorbance at 450 and 620 nm wavelength was measured using a Tecan M1000 microplate reader. Fitting curves were calculated using GraphPad Prism.

### Induction of costimulatory receptor expression on B-cells

PBMCs from three different donors were isolated from human buffy coat by Ficoll density gradient centrifugation and untouched B-cells were purified by negative magnetic enrichment using a B-cell isolation kit II (Miltenyi Biotec) according to the manufacturer’s instructions. 2×10^5^ B-cells in 100μl RPMI-1640 + 10% Human AB Serum were stimulated with CD40 antibody concentrations ranging from 500 to 0.2 ng/ml for 48h. Stimulated B-cells were harvested, stained using fluorophore-labeled antibodies against CD86 and CD80 and analyzed by flow cytometry on a BD FACSVerse device. Fitting curves were calculated using GraphPad Prism.

### Competition of CD40 antibodies with CD40L binding to cell-expressed CD40

HEK-Blue™-CD40L cells were preincubated with antibodies at their EC90 binding concentration for 30 minutes at 4°C. Recombinant CD40L protein was added at concentrations ranging from 10,000 to 9.8 ng/ml and cells were incubated for 60 minutes at 4°C. CD40L bound to cell-expressed CD40 was detected using secondary DyLight 405-conjugated anti-mouse IgG (Jackson Laboratories) while the anti-CD40 antibodies were detected using an Alexa Fluor 488-conjugated goat anti-human-F(ab)2 (Jackson Laboratories). Cells were analyzed using a FACSVerse instrument (BD).

### Competition of anti-CD40 antibodies for binding to human CD40

Antibodies were coated to 384-well Maxisorp plates at a concentration of 625 ng/ml in PBS for 60 minutes followed by a blocking step with PBS, 2% BSA, 0.05% Tween for 70 minutes. All antibodies were incubated separately at a concentration of 10 μg/ml in tubes together with 330 ng/ml HIS-tagged CD40 recombinant protein and 4 μg/ml peroxidase-coupled anti-HIS detection antibody (Sigma-Aldrich) for 60 minutes in ELISA buffer (PBS, 0.5% BSA, 0.05% Tween). The plate was washed three times with PBS, 0.1% Tween before the antibody/HIS-CD40/anti-HIS-peroxidase mixes were added to the wells of the plate. The plate was incubated for 60 minutes. Wells were washed six times with PBS, 0.1% Tween and 15μl/well TMB substrate solution (Invitrogen) was added. The reaction was stopped with 15μl/well Stop solution (1M HCl) and absorbance at 450 and 620 nm wavelength was measured using a Tecan M1000 microplate reader.

### Stem-cell-humanized mouse model

Animal work was performed in accordance with the guidelines of the National Institutes of Health and the legal requirements in Germany. Human stem cells were purified from umbilical cord blood with the written consent of patients and according to the ethical guidelines of the University of Erlangen.

Nod/Prkdcscid/IL2rg -/- FcRγ -/- (female and male mice) were generated by crossing the individual mouse strains, followed by backcrossing to the Nod/Scid background. Mice were irradiated sublethally with 1.4Gy within the first 24 hours after birth, followed by engraftment with 20,000-50,000 human hematopoietic stem cells isolated from umbilical cord blood (hCD34+ MACS). When the mice were aged 10-12 weeks the percentages of humanization in the peripheral blood were checked via FACS analysis. Successfully humanized mice (≥ 5% hCD45+ cells) were injected i.v. with different amounts of anti-CD40 antibody or isotype control antibody in 100 μl 1xPBS. Body weight and temperature were measured before treatment and at different time points after antibody injection.

### Analysis of human leucocytes in murine peripheral blood

To determine relative (or absolute) changes in white blood cell composition in the blood, FACS analyses were performed. Peripheral Blood (100-150 μl) was collected by retro-orbital or venal puncture. Erythrocytes were lysed by adding 900 μl of ddH2O for 20 sec. After the addition of 100 μl of 10x PBS the reaction was stopped, and the probes were centrifuged at 600xg, 5 min., RT. The cell pellet was resuspended in Fc-Block (0.5 μg/well, 2.4G2) and incubated for 15 min. at 4°C. After one centrifugation step (1,400 rpm., 5 min., 4°C) the cells were stained with fluorochrome-conjugated antibodies for FACS Analysis (all antibodies from Biolegend) for 15 min. at 4°C. For live/dead cell exclusion DAPI was added at dilution 1:5,000. After centrifugation (1,400 rpm., 5 min., 4°C) cells were resuspended in 100 μl FACS buffer and analyzed on a FACS Canto II.

### Serum cytokine analysis

Human cytokine levels in murine serum were detected by LEGENDplex™ Multi-Analyte Flow Assay Kit, an immunoassay based on beads. Peripheral blood was obtained as described above and centrifuged at 10,000xg for 5 min., afterwards serum was collected and stored at −80°C. Samples were diluted 1:2 with assay buffer and 12.5 μl of each of the following reagents was mixed in a 96-well-plate: assay buffer, sample or standard, mixed beads and detection antibody were added per well. The mixture was incubated at RT for 2h on a shaker (1000 rpm). 12.5 μl of SA-PE/well was added and again shaken (1,000 rpm) at RT for 30 min. After one washing step cells were resuspended in 200 μl wash buffer and analyzed on a FACS Canto II.

### Immunohistochemistry of liver and kidney

Liver and kidneys were frozen at −80°C in OCT, cut into 5 μM sections and fixed with acetone for 2.5 min. Sections were incubated in blocking solution (5% goat serum in PBS) for 1 hour at RT before fluorochrome-conjugated antibodies were added in 5% goat serum in PBS and incubated for 30 min. at RT in the dark. Slides were rinsed 3 times with 1xPBS, mounted with a drop of mounting medium and dried for 30 min. at RT in the dark. Stained sections of liver and kidney were analyzed on an Axiovert 200M.

### AST / ALT measurement

Concentrations of aspartate aminotransferase (AST) and alanine aminotransferase (ALT) in serum samples were determined using Alanine Aminotransferase (ALT) Activity Colorimetric Assay Kit and Aspartate Aminotransferase (AST) Activity Colorimetric Assay Kit from Biovision following the manufacturer’s instructions.

## Supplementary Materials

Fig. S1. Induction of NF-κB signaling by 303 humanized hIgG1-LALA CD40 antibodies in HEK-Blue-CD40L™ gene reporter cells.

Fig. S2. CD40 antibody binding to HEK-Blue-CD40L™ cells in the presence of CD40L

Fig. S3. Immune-histology analysis of anti-CD40-antibody-treated human-stem-cell-transplanted mice

Fig. S4. Analysis of ALT and AST plasma levels after anti-CD40 antibody treatment

Table S1. CDR sequence divergence of novel CD40 agonistic antibodies

Table S2. Survival of antibody-treated human-stem-cell-transplanted mice

## Acknowledgements

This work was funded by MAB Discovery GmbH. A patent application on humanized CD40 antibodies is pending. S.F. is founder and CEO of MAB Discovery GmbH.

## Author contributions

K.B., A.C., F.N., S.F. designed research and analyzed data; E.B., N.J., C.P., A.I., C.R. performed research; L.K., S.P. generated and produced CD40 antibodies; K.B., A.C., F.N., S.F. wrote the paper.

## Supplementary Material

**Suppl. Figure 1.**
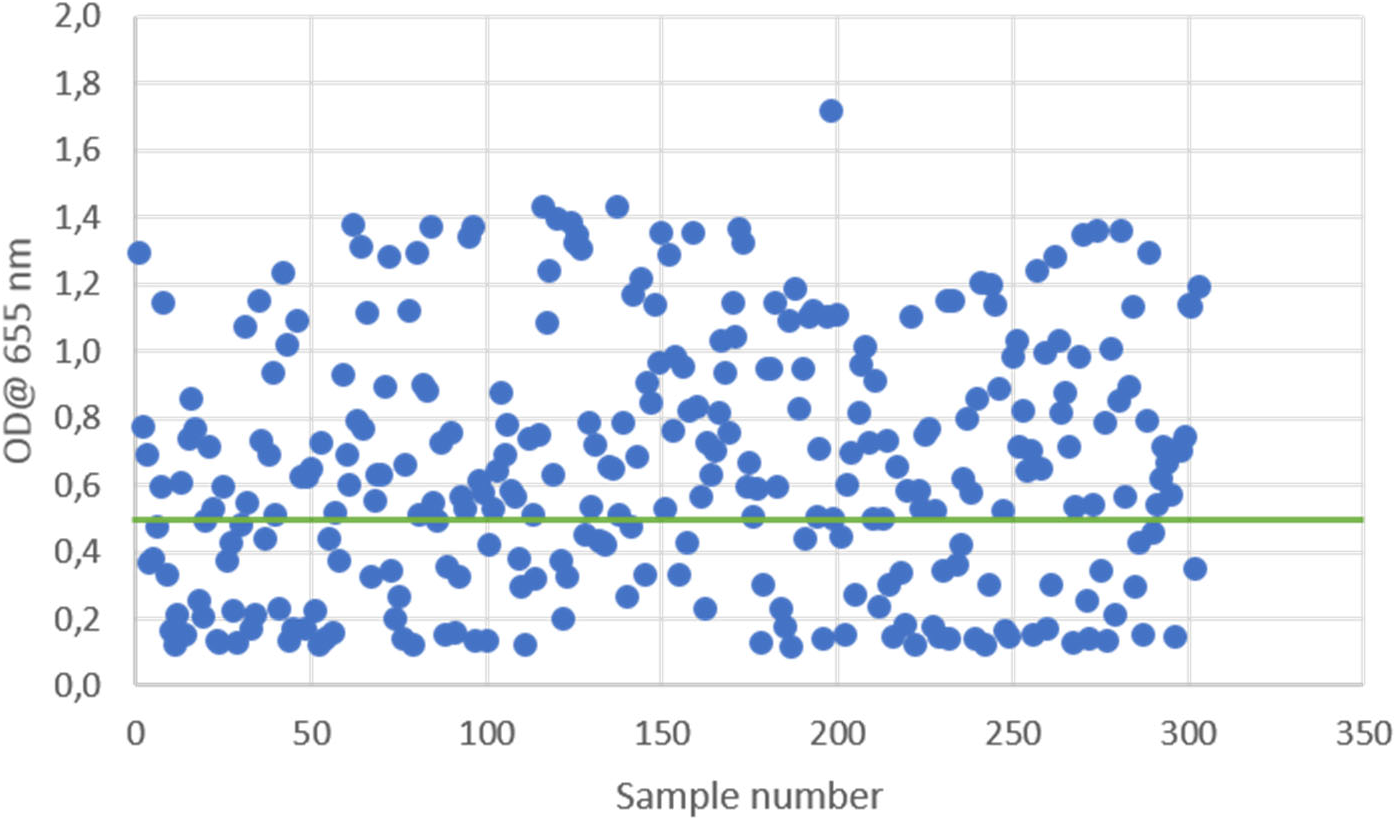
Induction of NF-κB signaling by 303 humanized hIgG1-LALA CD40 antibodies in HEK-Blue-CD40L™ gene reporter cells. Antibodies were incubated for 6 hours at a concentration range of 80 to 10000 ng/ml. Each dot indicates the maximum OD measured at 655 nm for each antibody. The green line indicates the threshold above which antibodies were classified as functional in this assay.

**Suppl. Figure 2.**
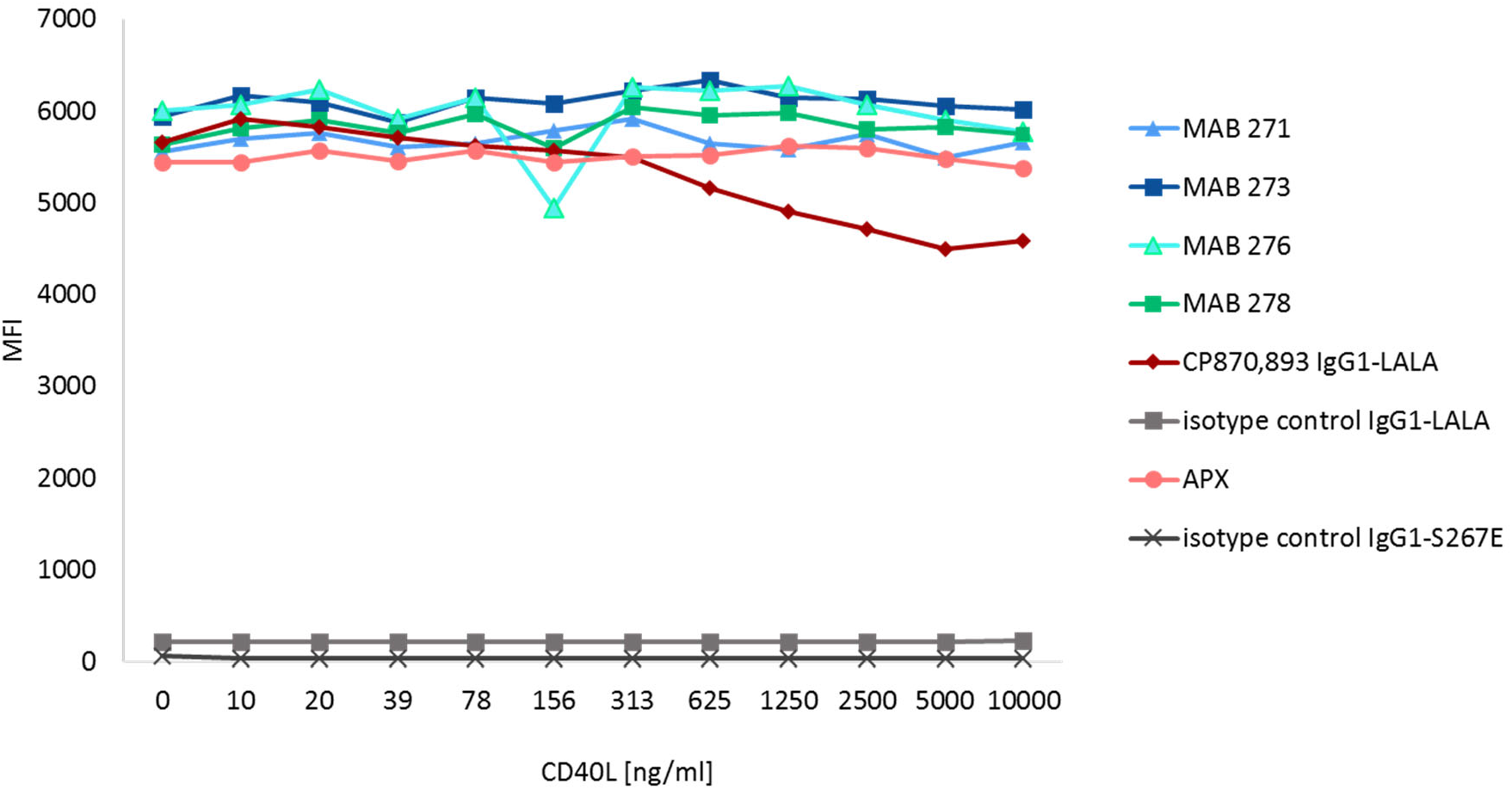
CD40 antibody binding to HEK-Blue-CD40L™ cells in the presence of CD40L. Cells preincubated with CD40 antibodies or isotype controls were incubated with increasing concentrations of mouse-Fc-tagged CD40L. Antibody binding was detected using a fluorophore coupled anti-human IgG and analyzed by flow cytometry.

**Suppl. Figure 3.**
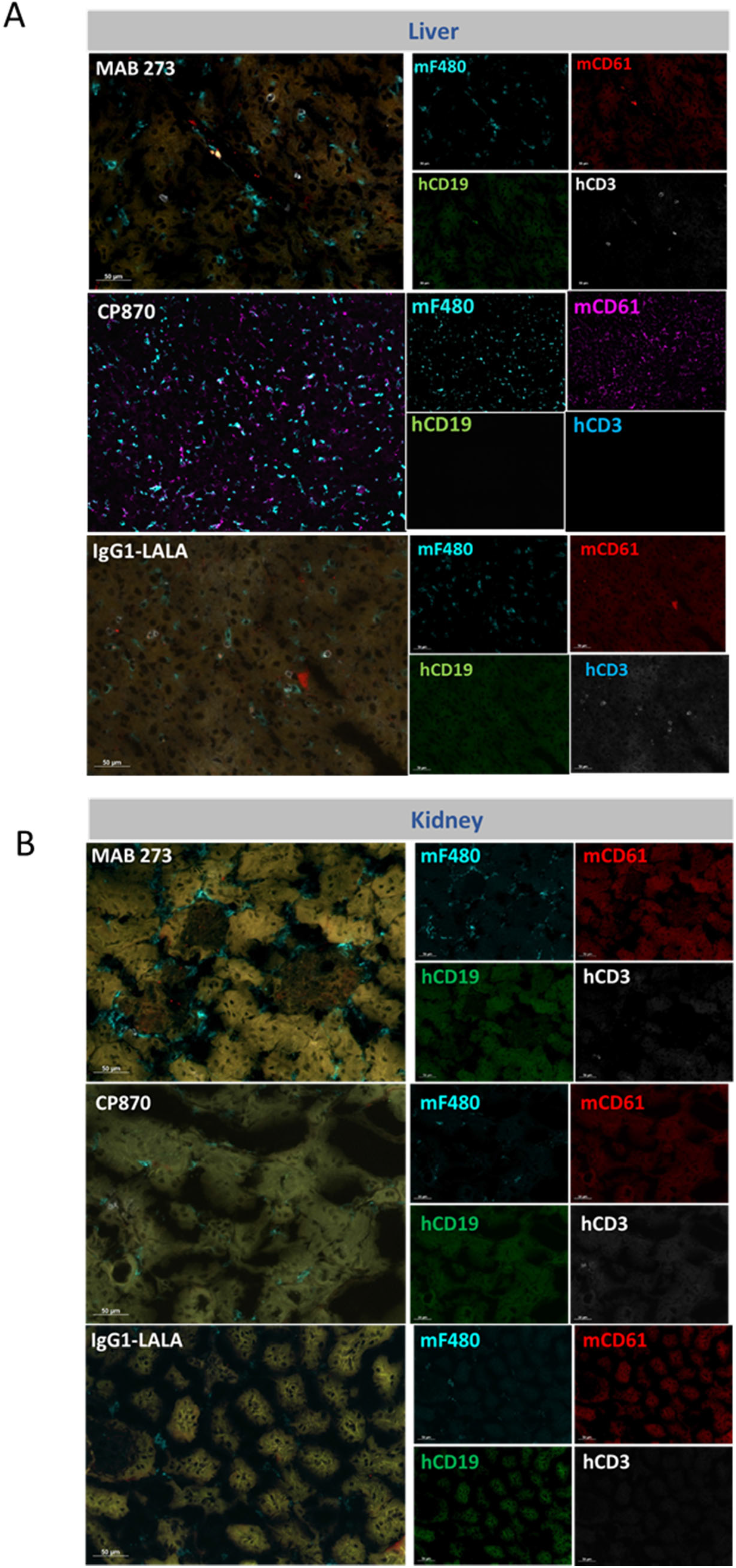
Immune-histology analysis of anti-CD40-antibody-treated human-stem-cell-transplanted mice. Shown is the analysis of liver (A) and kidney sections (B) of mice injected with MAB 273, CP870,893 or hIgG1-LALA control antibodies. Sections were stained for mouse macrophage-(mF4/80), mouse platelet-(mCD61), human B cell- (hCD19) and human T cell- (hCD3) markers. White arrows indicate hCD3 positive T-cells detected in liver sections of MAB 273 treated mice.

**Suppl. Figure 4.**
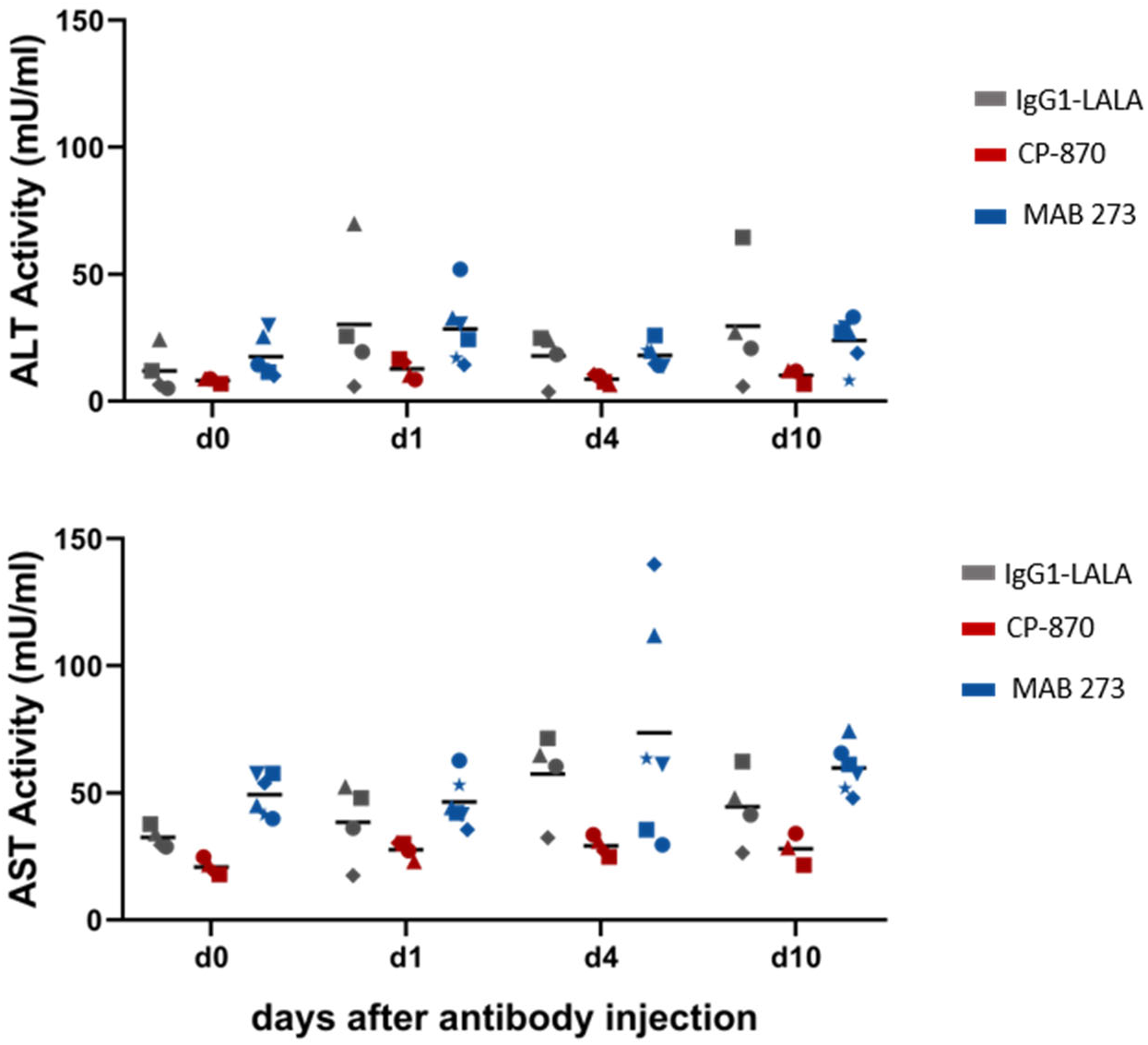
Analysis of ALT and AST plasma levels after anti-CD40 antibody treatment. Shown are the concentrations of the liver transamidases ALT and AST determined in the serum of human stem cell transplanted mice injected with the respective antibodies. Symbols indicate individual mice.

**Table S1:** CDR sequence divergence of novel CD40 agonistic antibodies. Differences of amino acids at defined positions are indicated as % difference for the heavy and light chain CDR sequences (HC/LC).

| Heavy chain CDRs |  |  |  | Light chain CDRs |  |  |  |
| --- | --- | --- | --- | --- | --- | --- | --- |
|  | MAB 273 | MAB 276 | MAB 278 |  | MAB 273 | MAB 276 | MAB 278 |
| **MAB 271** | 20% | 37% | 56% | **MAB 271** | 48% | 53% | 48% |
| **MAB 273** | - | 30% | 50% | **MAB 273** | - | 56% | 52% |
| **MAB 276** | - | - | 58% | **MAB 276** | - | - | 53% |

**Table S2:** Survival of antibody treated human stem cell transplanted mice. The table represents the total number of treated human stem cell transplanted mice from all experiments and the number of euthanized mice per treatment group.

| Treatment | Total number of mice | Number of euthanized mice |
| --- | --- | --- |
| **MAB 273** | **9** | **0** |
| **CP-870,893** | **13** | **5** |
| **IgG1-LALA** | **8** | **0** |

